# Calcium transients in nNOS neurons underlie distinct phases of the neurovascular response to barrel cortex activation in awake mice

**DOI:** 10.1101/2022.10.03.510654

**Authors:** Sung Ji Ahn, Antoine Anfray, Josef Anrather, Costantino Iadecola

**Author notes:** **Corresponding authors:** Costantino Iadecola, MD, Sung Ji Ahn, PhD, Feil Family Brain and Mind Research Institute, Weill Cornell Medicine, 407 East 61st Street, RR-303, New York, NY 10065.

## Abstract

Neuronal nitric oxide (NO) synthase (nNOS), a Ca^2+^ dependent enzyme, is expressed by distinct populations of neocortical neurons. Although neuronal NO is well known to contribute to the blood flow increase evoked by neural activity, the relationships between nNOS neurons activity and vascular responses in the awake state remain unclear. We imaged the barrel cortex in awake, head-fixed mice through a chronically implanted cranial window. The Ca^2+^ indicator GCaMP7f was expressed selectively in nNOS neurons using adenoviral gene transfer in nNOS^cre^ mice. Air-puffs directed at the contralateral whiskers or spontaneous motion induced Ca^2+^ transients in 30.2±2.2% or 51.6±3.3% of nNOS neurons, respectively, and evoked local arteriolar dilation. The greatest dilatation (14.8±1.1%) occurred when whisking and motion occurred simultaneously. Ca^2+^ transients in individual nNOS neurons and local arteriolar dilation showed various degrees of correlation, which was strongest when the activity of whole nNOS neuron ensemble was examined. We also found that some nNOS neurons became active immediately prior to arteriolar dilation, while others were activated gradually after arteriolar dilatation. Discrete nNOS neuron subsets may contribute either to the initiation or to the maintenance of the vascular response, suggesting a previously unappreciated temporal specificity to the role of NO in neurovascular coupling.

## Introduction

The neurovasculature is critically important for the maintenance of brain health ^1^. One of its vital functions is to preserve the metabolic homeostasis of the brain by matching energy substrates delivery and waste removal with the degree of metabolic activity of brain cells ^2^. This phenomenon termed functional hyperemia results from the concerted action of a wide variety of cell types at different levels of the cerebral vasculature ^2^. Accumulating evidence suggests that functional hyperemia is initiated by neural signals at the site of activation leading to endothelial hyperpolarization in local microvessels ^3-5^. Such hyperpolarization, in turn, is transmitted retrogradely through inter-endothelial and myo-endothelial junctions to hyperpolarize upstream contractile mural cells resulting in vasodilatation and increased flow to the activated area ^4-6^. The spatiotemporal correspondence between neural activity and functional hyperemia is at the bases of functional MRI signals widely used to localize brain function ^7, 8^. However, the nature of the neuronal signals and their relationships to the hemodynamic response remain poorly understood.

Numerous molecular signals have been implicated in functional hyperemia ^9-12^. However, nitric oxide (NO), a potent vasodilator released by a subpopulation of GABAergic neurons expressing the neuronal isoform of NO synthase (nNOS or *Nos1*)^13-16^, is widely recognized as a major contributor^9, 12, 17^. nNOS neurons fall into distinct types based on their histochemical features ^13^. Type 1 neurons have long range projections reaching the contralateral hemisphere and have been linked to sleep and wakefulness ^18-20^. Type 2 neurons have the morphological features of interneurons with short bipolar or multipolar projections ^13^. Both types also express mainly neuropeptide Y, somatostatin, and/or vasoactive intestinal peptide, have frequent contacts with blood vessels and have been implicated in neurovascular regulation ^13, 16, 21-24^. Thus, the increase in cerebral blood flow (CBF) elicited by brain activation is associated with NO release ^25-27^, while pharmacological inhibition or genetic deletion of nNOS, as well as depletion of nNOS neurons attenuates the increase in CBF produced by neural activity in different brain regions ^27-33^, a finding also observed in humans ^34^. Furthermore, chemical, optogenetic or chemogenetic activation of nNOS neurons increases local CBF ^25, 35-38^, attesting to their potential for neurovascular regulation. Despite these advances, the mode of activation of the nNOS neural network, and the spatiotemporal relationships between nNOS neuron activity and arteriolar dilatation have not been established.

Most studies investigating the link between neural activity and CBF have been performed in anesthetized animals. While unavoidable and necessary in certain experimental settings, anesthesia has potent neuronal and vascular effects that alter the neurovascular mechanisms underlying functional hyperemia ^39-41^. This problem is particularly concerning when attempting to relate nNOS neural network activity with vascular responses since anesthetics target differently GABAergic and glutamatergic neuronal populations, altering network activity and excitatory-inhibitory balance ^42^. Consequently, the neurovascular mechanisms driving functional hyperemia in the anesthetized state may not accurately reflect those operating in the awake state ^43^.

In the present study we used 2-photon excited fluorescence microscopy (2PEF) in awake mice to investigate the relationship between nNOS neuronal network activity, by the Ca^2+^ indicator GCaMP7f, and functional hyperemia in the whisker barrel cortex. We found that nNOS neurons exhibit diverse patterns of activity corresponding to spontaneous or evoked activation of the whisker barrel cortex or grooming-related motion. Correlation with the resulting arteriolar dilatation unveiled early and late patterns of nNOS neuron activity suggesting distinct roles in the initiation and maintenance of the hemodynamic response. Surprisingly, no link was observed between the distance of active nNOS neurons from nearby arterioles and their involvement in arteriolar dilatation. The data provide a first insight into the spatiotemporal dynamics of the events linking nNOS neuron activity to microvascular function and may aid in the interpretation of the alterations in neurovascular coupling occurring in pathological states in which nNOS neurons are compromised ^44-46^.

## Materials and Methods

### Animals

All procedures were approved by the Institutional Animal Care and Use Committee of Weill Cornell Medicine (protocol number 0711-687A) and performed in strict accordance with ARRIVE guidelines ^47^. Studies were performed in 3-4 months old mice of both sexes on C57BL/6J genetic background. NOS1^cre^ breeders were purchased from the Jackson Laboratory (#017526) and breed in house with mice on C57BL/6J genetic background.

### Animal preparation

Optical access to brain was achieved through a long-term implanted glass-covered cranial window, as described previously ^48^. Briefly, animals were anesthetized using isoflurane (1.5–2% in oxygen) and placed on a feedback-controlled heating blanket that maintained body temperature at 37°C (50-7053P; Harvard Apparatus). After removing the scalp and periosteum from the skull surface, a custom-made titanium head post was glued onto the right hemisphere over the somatosensory cortex using dental cement (C&B Metabond, Parkell). A 3-mm diameter bilateral craniotomy was performed over parietal cortex using a dental drill. The exposed brain was covered with sterile saline and sealed with a 3-mm square borosilicate coverslip glass (CS-3S, Warner Instruments) using cyanoacrylate glue and tissue adhesive (1469SB; 3M). For nNOS neuron specific GCaMP7f expression, AAV PHP.eB Syn-FLEX-GCaMP7f (104492-PHPeB, Addgene) was diluted (4 × 10^10^ vg / 100 μl sterile saline) and injected retroorbitally. All mice were allowed to recover for at least 7 days before handling and habituation to the head restraint for awake *in vivo* imaging. Mice were gradually habituated to longer periods (1, 5, 10 up to 40 min) to the head-restraint under the microscope. All mice were rewarded with sweetened milk. In some mice the femoral artery was cannulated under isoflurane anesthesia for recording of arterial pressure and blood gas while head restrained. As in previous studies ^49^, wounds were treated with lidocaine ointment and mice remained calm after awakening. Mice had stable blood pressure (mean arterial pressure: 84.8±2.5 mmHg) and blood gases (pH 7.329±0.016, PaCO_2_ 35.2±1.3 mmHg, PaO_2_ 116.8±3.9 mmHg, O_2_ saturation 98.3±0.2 %).

### Two-photon imaging in awake mice

As described in detail elsewhere ^44, 48^, to visualize the microvasculature mice were briefly anesthetized with isoflurane (1.5–2%, during ∼2 min) and injected retroorbitally with 70 kDa Texas Red-conjugated dextran to label the plasma (50 μL, 2.5% w/v, Invitrogen). Mice were allowed to recover fully prior to awake imaging. For 2PEF imaging, mice were head fixed using a titanium headplate and placed inside a falcon tube on a soft surface. Imaging was performed on a Fluoview 2-photon microscope (FVMPE; Olympus) and excitation pulses were provided by a solid-state laser (InSight DS+; Spectraphysics). For acquiring two channel movies (GCaMP7f and Texas Red), the excitation laser was tuned to 910 nm and fed through a primary dichroic mirror (FV30-NDM690). Emission light went through an IR blocking filter (FV30-BA685RXD) and then guided toward gallium arsenide phosphide (GaAsP) photomultiplier tubes (PMTs) by FV30-SDM-M mirror. GCaMP7f and Texas Red fluorescence were collected using an FV30-FGR filter cube (green: 495-540 nm, red 575-645 nm, separated by 570 nm dichroic mirror) mounted in front of GaAsP-PMTs. Image stacks and movies were acquired through Fluoview software (FV31S-SW, version 2.3.1.163; Olympus). First, a map of the vasculature was taken through a ×5 objective (MPlan N 5□×□0.1 NA, Olympus) and compared with laser speckle map to locate the same imaging field in the whisker barrel cortex and identify pial vessels branches that feed the barrel cortex. Once the blood vessels to be imaged were identified, we switched to a ×25 objective (XLPlan N 25□×□1.05 NA, Olympus). Two channel movies of 509 × 509 μm^2^ (512 × 512 pixel) were acquired using a resonant scanner with a framerate of 3.8 Hz (4 lines averaged, 5 minutes long). During imaging, mice received trains of nitrogen 5 sec or 30 sec long (air-puffs) angled toward the tip of whiskers contralateral to the imaged cortex. We used lowest setting from a picospritzer to generate gentle light air-puff (approx. 10 psi, Picospritzer II, Parker). The pressure of the air puff was adjusted so that only the whiskers were displaced, but the mouse was not startled and did not display any signs of agitation or discomfort (Supplementary movie 1). An optogenetic TTL pulse generator (OTPG_4, Doric Lenses) was triggered directly from the Olympus breakout box at the start of data acquisition and a Doric GUI platform allowed us to program duration and frequency of air puff train. Mice were continuously monitored during experiments through live videos with an infrared camera (120 frames/second) illuminated with 850 nm LED via Bandicam software and focused on the whiskers that received the air-puffs. We placed a 850 nm short pass filter (FES0850, Thorlabs) in front of the lens of the webcam to block the excitation laser light. The live movie during each image acquisition was saved for off-line analysis of whisking and grooming related motion (Supplementary movie 2). We programmed an Arduino board to signal Bandicam software to synchronize the video recording with the imaging acquisition.

### Analysis of *in vivo* calcium transients

The GCaMP7f channel of the two-channel movie was motion corrected to account for small xy displacement using the motion correction module of EZcalcium (https://github.com/porteralab/EZcalcium), which implemented the Non-Rigid Motion Correction (NoRMcorre) strategy ^50^. Then soma of nNOS neuron were visually inspected and manually contoured on CalciumDX software ^51^ (https://github.com/ackman678/CalciumDX). Signal intensity from each soma contour over each movie frame was saved for further analysis using custom written Matlab code. Fluorescence signal time series (ΔF/F: change in fluorescence divided by baseline fluorescence) were calculated for each individual neuron. For 5 seconds whisker air puffs, the mean of the lower 25% in a 15-second sliding window was used to calculate the baseline fluorescence for each cell to account for both differences in GCaMP expression and de-trending for slow time-scale changes in fluorescence ^52^. For analysis of 30 sec whisker air puffs we used the mean of the lower 10% in 50-second sliding window as baseline fluorescence. To determine the onset and termination of a calcium transient, the ΔF/F signal was processed through a 4^th^ order zero phase low pass filter (Matlab functions: butter, filtfilt) and marked active when ΔF/F exceeded two standard deviations above the baseline fluorescence. The termination of a calcium transient was identified as occurring when ΔF/F fell below 0.5 standard deviation (SD) above the baseline fluorescence, SD was determined from the entire population in each experiment to identify active cells not confounded from background noise. To accurately detect firing onset accounting for the time it takes to reach two SD of ΔF/F, we considered activation time when the ΔF/F derivative increased more than a half SD. Neuronal ensemble activity over time was simply calculated as the percent of cells active in each frame. Statistically significant multicellular ensemble events were determined when the percent of simultaneously active cells exceed the percent expected by chance given the observed level of activity in the field of view. The threshold was calculated as detailed elsewhere^52^. Cells that did not exhibit any statistically significant Ca^2+^ transients during a given recording session were excluded from analyses.

### Analysis of penetrating vessel diameter

The Texas red channel of the two-channel movie was used for diameter analysis. Vessels cross sections with a diameter > 10 μm were examined with ‘plot Z axis profile’ in ImageJ. Arterioles and venules were identified based on vasoactivity and direction of flow. To estimate diameter of a penetrating arteriole accurately, we used thresholding in Radon space (TiRS) (https://github.com/DrewLab/Thresholding_in_Radon_Space) ^53^. This TiRS method is robust to noise and shape changes when arteriole dilate / constrict and accurately computes cross sectional area of a penetrating arteriole. Arteriolar diameter in time series were processed through a 2^nd^ order zero phase low pass filter (Matlab functions: butter, filtfilt) before further analysis. The percent diameter change due to motion and short air-puff (5 seconds) was calculated from baseline diameter right before dilation and peak dilation. Percent diameter change during 30 seconds air-puff was calculated by averaging 5 seconds at the baseline and diameter at 25-30 seconds during air-puff.

### Analysis of whisking and motion

Time locked videos from the mouse infrared camera during image acquisition were analyzed to quantify whisking and grooming motion. For whisker analysis, we draw a box encompassing the whiskers and computed mean whisker angle by applying the Radon transform (Matlab function: radon) to each frame as previously described ^54^. The mean whisker position was low pass filtered to 30 Hz (Matlab functions: butter, filtfilt) then whisker acceleration was computed from the second derivative. The absolute whisker acceleration was smoothened and binarized by manual thresholding, empirically determined for each animal. To detect motion during grooming, we draw a box encompassing the mouse’s upper torso and forelimbs. We then calculated the mean difference of each pixel value between two consecutive frames, which quantified body position changes between each movie frame. The motion estimate was analyzed further as described for the whiskers above. We binarized both whisking and motion as changes in values between each frames in a 120 frames/second.

### Immunohistochemistry

Immunostaining of brain was performed as described before with some modifications ^48, 55^. Animals were anesthetized with sodium pentobarbital and transcardially perfused with 15 ml of phosphate buffered saline (PBS) (Sigma-Aldrich) followed by 30 ml of 4% w/v paraformaldehyde (PFA) (Fischer Scientific) in PBS. Brains were extracted, stored overnight in in 4% PFA in PBS at 4°C, then in 15% w/v sucrose in PBS for 8 hr at 4°C, followed by 24 hr in 30% sucrose, and then in 60% sucrose until sectioning. The tissue was then frozen and cut into 20-μm thick coronal sections on a cryostat, mounted onto microscope slides and kept at −80 °C. Brain sections were permeabilized with 0.5% Triton X-100 (Sigma) in PBS (PBST) for 30 min, blocked with 10% normal donkey serum (NDS) in 0.1% PBST for 30 min and incubated in 5% NDS-0.1% PBST at 4 °C overnight with the following primary antibodies against NOS1, GFP, GAD67, or CaMKII. The next day, after three washes with PBS, specimens were incubated with anti-chicken Alexa flour 488-, anti-rabbit Cy3- or anti-mouse Alexa flour 647-conjugated donkey secondary antibodies (1:200; Jackson ImmunoResearch Laboratories). Then, sections were washed with PBS and mounted using an anti-fade mounting medium that contained DAPI (Vetashield, H-1200, Vector Laboratories). Fluorescent images were collected using a confocal microscope (Leica TCS SP8) with a 20x/0.75 NA objective (11506517, Leica). The following primary antibodies were used: rabbit anti-NOS1 polyclonal antibody (1:300, AB5380, Sigma-Aldrich), chicken anti-GFP polyclonal antibody (1:500, A10262, Invitrogen), mouse anti-GAD67 monoclonal antibody (1:300, MAB5406, Sigma-Aldrich), rabbit anti-CaMKII monoclonal antibody (1:300, ab52476, abcam).

### Analysis of mRNA data

Mouse brain single cell mRNA data was downloaded from Mouse Brain Atlas (https://storage.googleapis.com/linnarsson-lab-loom/dev_all.loom) ^56^. We used Seurat (Vers. 4.1.0) ^57^ in the R statistical environment for analysis and data visualization. The expression matrix was downsampled to limit the number of cells in each cluster defined by “ClusterName” to 2000. The matrix was further reduced by rejecting cells with <500 or >30.000 UMI counts, or >5% mitochondrial genes. Log normalization, variable gene detection, scaling, and principal component analysis was performed in Seurat with default settings. UMAP was computed on the first 40 principal components. We then selected telencephalic neurons matching the occurrence of “TE” in the ClusterName metadata column and further rejected cells that originated from animals <28 days of age. UMAP plots were visualized with the *FeaturePlot* function in Seurat. Color scales are log-transformed normalized expression values.

### Correlation analysis

Neuronal ΔF/F for each neuron and determination of active/inactive state during time series are detailed above. We used Spearman correlation to correlate the activity (ΔF/F) between individual neurons and every other neuron over a 5 minute period where calcium transients (ΔF/F) values during inactive state were set to zero (**Figure 2F and G**). When matching the time each neuron fired during certain behavior such as whisking or motion (**Figure 2H**), we binarized neuronal activity as active or inactive, and correlated it to the binarized motion or whisking traces. Pearson correlations were calculated between neuronal calcium transients (ΔF/F) and arteriolar diameter (μm) within the same recording session over a 5 minute period. Calcium transients (ΔF/F) values during inactive state were set to zero (**Figure 3D**). We used normalized diameter change over time (0 – 1 range, Matlab function: normalize, ‘range’) and aligned the start of neuronal activity with the vascular response to account for the delay of arteriolar dilatation slightly (approx. 1 second). To this end, we found the delay between normalized arteriolar diameter change and normalized neuronal ensemble using (Matlab function: finddelay) and shuffled the data accordingly (Matlab function: circshift). For each neuron we selected the 30 seconds air-puff period with the least motion and calculated the Person correlation between ΔF/F and normalized arteriolar diameter to cover 5 seconds pre air-puff, and 5 seconds post. To quantify the delay between calcium transients (ΔF/F) and arteriolar dilation we run cross correlation (Matlab function: xcorr) of ΔF/F and normalized arteriolar diameter to determine the time lag of the two signals (Figure 4D).

### Image, data, and statistical analysis

All 2PEF images were analyzed using custom written code on Matlab (R2018a, R2021b, R2022a, Mathworks) unless otherwise noted (R2014b for CalciumDX, R2018a and above for EZCalcium). Immunohistological images are maximum z projections and representative 2PEF images are average projections in time. Representative correlation matrix on Figure 2F was plotted after sorting neuronal calcium transients in ascending order based on the values in the time when highest neuronal ensemble was observed to cluster/highlight neurons with functional connectivity. Statistical analyses were done using Matlab (R2021b, MathWorks) or Prism (v.9.4.0, GraphPad Software). Graphs in all plots represents the mean and error bars represent the SD. Data were tested for normal distribution by the D’Agostino-Person test before running any comparisons. When comparing among different animal, we used nested one-way ANOVA. To evaluate statistical differences between the linear correlations of two data sets, we generated one-way analysis of covariance models (Matlab function: aoctool) and ran multiple comparison test (Tukey-Kramer) of the group means (Matlab function: multcompare). To estimate the linear relationship between distance and correlation coefficient we used simple linear regression on Prism. The data that support the findings of this study and codes to plot and analyze data are available from the author on reasonable request.

## Results

### GCaMP7f is selectively expressed in nNOS neurons which include both excitatory and inhibitory subtypes

First, we sought to characterize the nNOS population expressing GCaMP7f. To achieve selective GCaMP7f expression in nNOS neurons we administered a GCaMP7f expressing adeno-associated virus driven by the synapsin promoter and containing a *Cre* responsive flip-excision switch (AAV PHP.eB Syn-FLEX-GCaMP7f, i.v.) in a nNOS *Cre* driver mouse line (NOS1^cre^). The GCaMP7f AAV transduced nNOS neurons with high efficiency. Immunohistochemical analysis showed that 95.3 ± 2.8 % (mean ± SD) express both nNOS-immunoreactivity and GCaMP7f (**Figure 1A, C**). Of the GCaMP7f expressing neurons, 87.2 ± 3.2 % were inhibitory (GAD67 positive) and 12.8 ± 3.2 % were excitatory (CaMKII positive) (**Figure 1B, C**). To confirm that both excitatory and inhibitory neurons expressed nNOS, we mined a mouse single-cell RNAseq dataset ^56^. We found that some nNOS neurons express the inhibitory neuron transcript *Gad1* (87.0 %) and others the excitatory neuron transcript *Slc17a7* (11.4 %) (**Figure 1D**). Therefore, using this viral gene transfer approach we were able to induce GCaMP7f expression in the majority of nNOS neurons, both excitatory and inhibitory.

**Figure 1.**
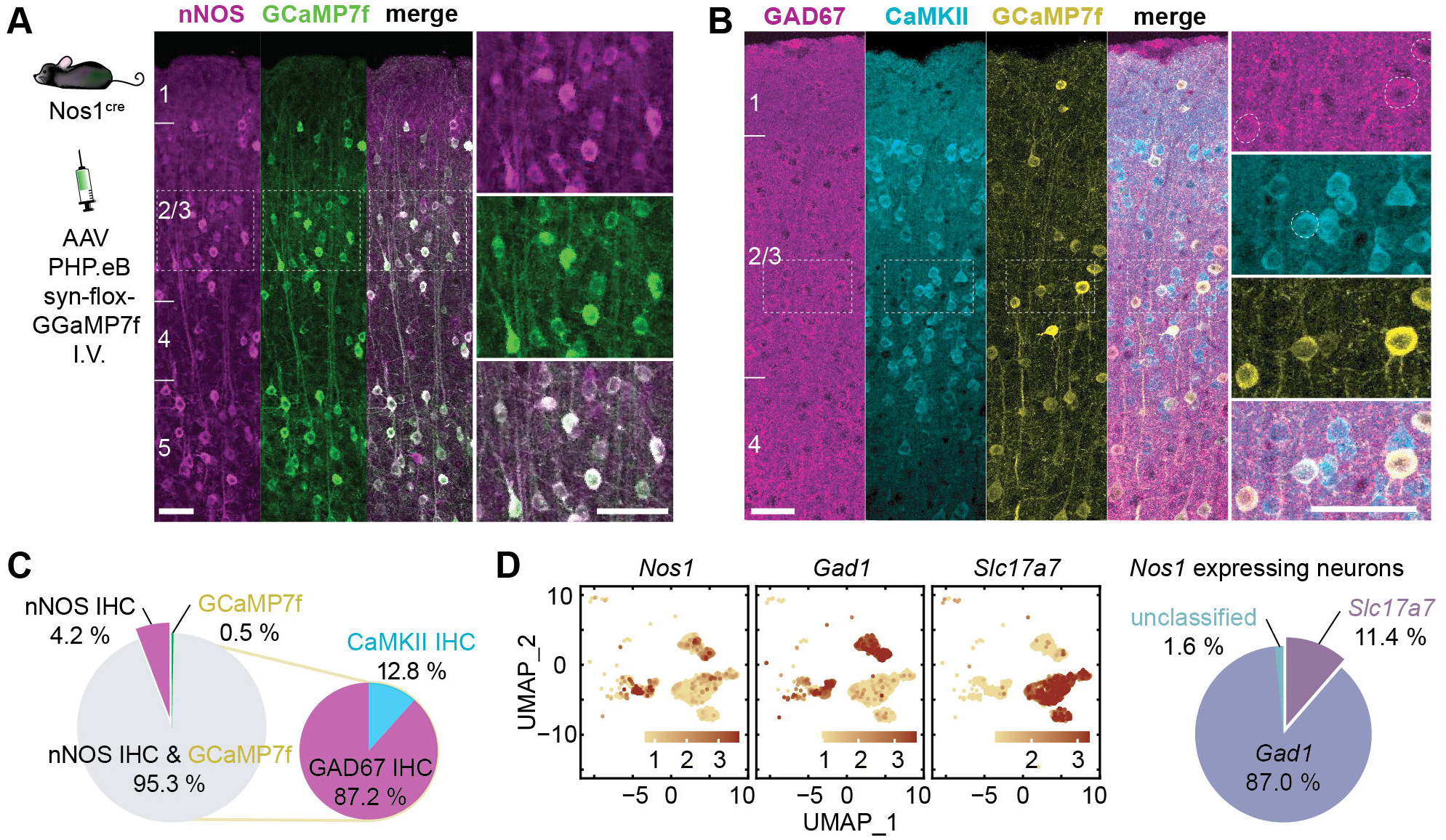
Viral gene transfer-induced expression of GCaMP7f in nNOS neurons of NOS1^cre^ mice. **A**: GCaMP7f was expressed in NOS1^cre^ mice using systemic administration of an AAV-PHP.eB viral vector. The localization of GCaMP7f expression in nNOS neurons in the cortical layers was verified by immunohistochemistry (IHC). Right panels are enlargements of the dashed squares in the left panel (n=2,744 neurons in 5 mice). **B**: GCaMP7f expression in inhibitory (GAD67) or excitatory (CaMKII) neurons of nNOS^cre^ mice treated with the AAV-PHP.eB viral vector. **C**: Neurons (%) expressing nNOS, GCaMP7f or both, and, among GCaMP7f positive neurons, those expressing GAD67 or CaMKII (n=690 neurons in 3 mice). **D**: Single-cell RNAseq data (mousebrain.org/development/downloads.html) showing expression of *Nos1* both in excitatory (*Slc17a7*) and inhibitory (*Gad1*) neurons (%), confirming the finding by immunohistochemistry (**B**). Scale bar=50μm throughout.

### nNOS neurons differentially respond to whisker activity and motion

We then performed imaging of GCaMP7f labeled nNOS population in the whisker barrel cortex in head-restrained awake mice through a chronic cranial window (**Figure 2A**). To localize the whisker barrel cortex, we used laser speckle imaging during mechanical stimulation of the contralateral whiskers, as previously described ^44^. The laser speckle signal increased about 20% in the whisker barrel cortex (**Figure 2B**). Awake mice were then head fixed under the two-photon microscope and, using the laser speckle map as a guide, we imaged the whisker barrel cortex during delivery of air puffs to the contralateral whiskers. Although head-restrained, the mice were free to readjust their body position and from time to time exhibited grooming behavior (Supplementary movie 2). We defined ‘motion’ as spontaneous movements that involved grooming or changes in posture. ‘Whisking’ includes both air-puff induced whisker deflection and spontaneous whisking. This experimental setting allowed us to map the spatiotemporal dynamics of cortical nNOS neurons activity in behaving mice with and without whisker air-puffs (**Figure 2D**). Analysis of the Ca^2+^ signal revealed that nNOS neurons exhibited different patterns of activity. Some neurons responded faithfully to all the whisker puffs, but not to spontaneous whisking or motion (**Figure 2D;** neuron 3). Other neurons responded to spontaneous or evoked whisking (neuron 1) and some predominantly to motion (neuron 2).

**Figure 2.**
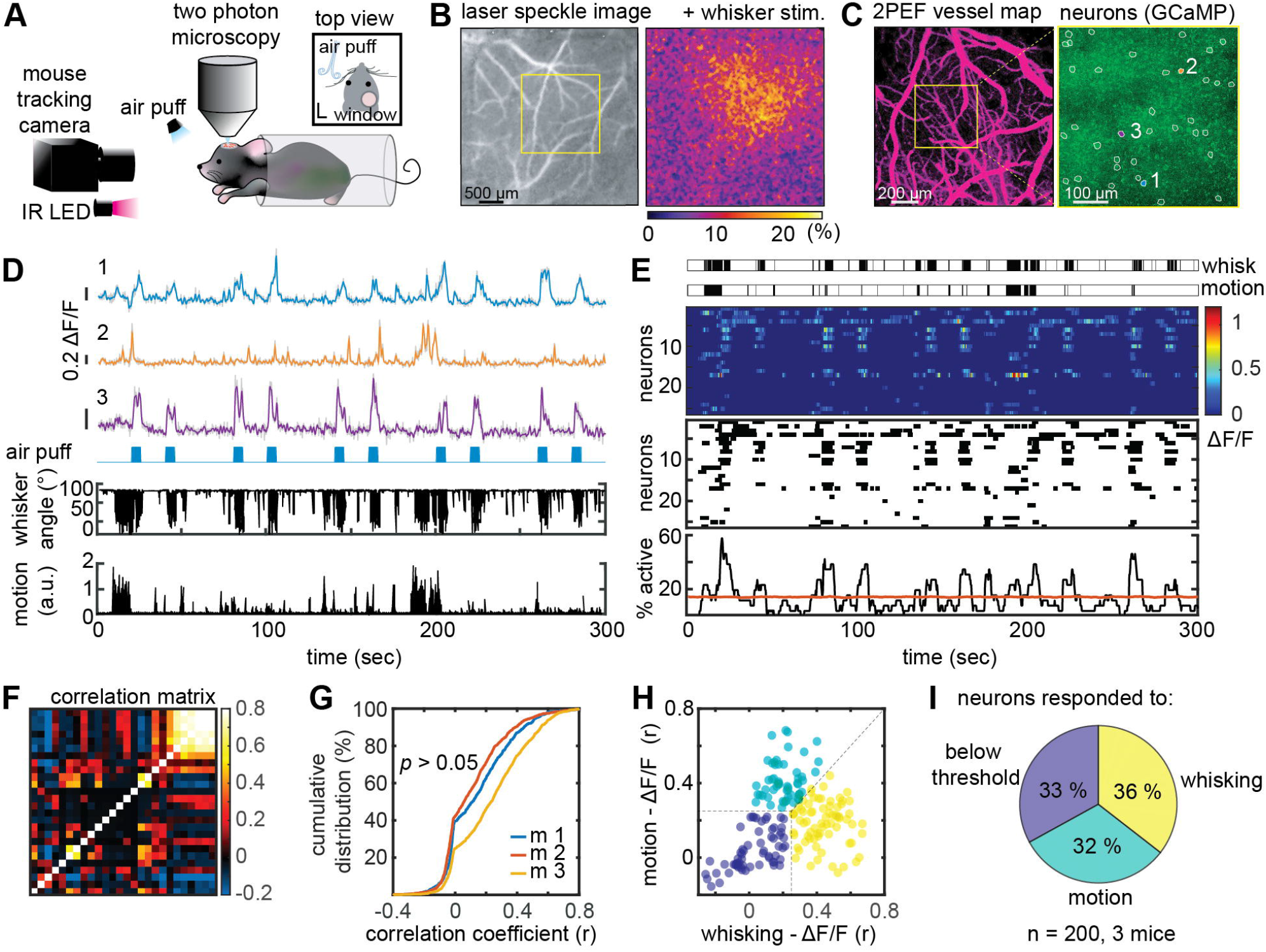
*in vivo* imaging of nNOS neurons in awake, head-fixed mice during air-puff whisker stimulation. **A**: Experimental setup for 2PEF imaging during whisker air-puff stimulation, which includes an infrared (IR) camera for tracking whisker and upper limb movements. **B**: Localization of the hemodynamic response evoked by whisker stimulation in the contralateral whisker barrel cortex by laser speckle imaging. **C**: <u>Left panel</u>: 2PEF map of pial vessels (within the yellow square in **B**) imaged at 5x. <u>Right panel</u>: GCaMP7f expressing nNOS neurons in layer 2/3 (outlined in white) imaged at 25x. The Ca^2+^ activity of neurons labelled 1, 2 and 3 is shown in **D. D**: Ca^2+^ activity (ΔF/F see methods) in neurons 1, 2 and 3 during 10 air puffs each lasting 5 sec (blue marks). Whisker displacement and limb motion are also displayed. Notice that neuron 3 responds to air puffs, while neuron 2 mainly to motion. Neuron 1 exhibits mixed responses. **E**: Representative raster plots of ΔF/F (top), neuronal activity (middle), and proportion of simultaneously active cells over time (neuronal ensemble, bottom) from the neuronal population shown in (**C**, right panel). Each row represents a single cell. Ensemble activity above the orange line is statistically significant. **F**: Correlation matrix quantifying functional connectivity between each neuron and every other neuron shows a broad range of correlations (Spearman’s rho). **G**: Cumulative distribution of correlation coefficients across all animals shows that all mice had similar distribution (nested one-way ANOVA). **H**: scatter plot of the correlation coefficient between each neuron ΔF/F and whisker activity (x-axis) or motion (y-axis) showing three populations of nNOS neurons each responding predominantly to whisking, motion or neither. (**I**) Pie chart summarizing data in **H** from all mice. The dotted lines in **H** were used as cutoff for the pie chart.

We then examined the ensemble activity of the nNOS neurons evoked by air puffs or motion. We found a significant synchrony of Ca^2+^ activity with up to 58% of nNOS neurons firing at the same time (**Figure 2E**). To gain insight into the functional connectivity among nNOS neurons, we assembled a correlation matrix relating Ca^2+^ activity across all cells in a field of view (**Figure 2F**). We found a broad distribution of correlation coefficients with some neurons exhibiting synchronous Ca^2+^ activity (r: 0.5-0.8), and other less synchronous activity (**Figure 2G**). We then sought to provide additional evidence that specific subgroups of nNOS neurons responded to a specific stimulus, as suggested by inspection of the patterns of neural activity. Using an arbitrary correlation coefficient cutoff (r: 0.25), we found that some neurons were more responsive to whisking activity (36%) and others to motion (32%). Other neurons (33%) were weakly correlated (r <0.25) to whisking or motion (**Figure 2H, I**).

### nNOS neurons ensemble activity is strongly correlated with arteriolar dilatations

Since nNOS neurons have been linked to functional hyperemia ^25-33^, we next sought to examine the relationship between nNOS neuron Ca^2+^ activity and local arteriolar dilatation (**Figure 3A, B**). Arteriolar dilatations ranging from 3.4-22.0% promptly followed peaks of neuronal ensemble activity induced by air puffs, spontaneous whisking or motion (**Figure 3B, C**). Larger arteriolar responses were observed when air puffs and motion occurred at the same time (14.8±4.4%; p<0.05 from whisking + motion) (**Figure 3B, C**). Such larger hemodynamic responses were associated with greater ensemble activity resulting in a steeper linear relationship between the two (**Figure 3C**). However, the increased hemodynamic response could not be explained solely on the bases of increased neuronal activity since the vasodilatation was greater than that evoked by whisker stimulation alone with comparable ensemble activity (% of active neurons) (**Figure 3C**). Therefore, we examined if changes in blood pressure could contribute to larger hemodynamic response, in separate awake mice (n=5) equipped with an indwelling femoral artery catheter we monitored blood pressure during whisker stimulation and motion. We found that mean arterial pressure remained stable during 30 sec whisker stimulation (baseline: 84.8±2.5 mmHg; whisker stimulation: 84.7±4.0 mmHg; p>0.05), but was transiently increased during spontaneous motion (baseline: 84.8±2.4 mmHg; motion: 97.7 ± 4.0 mmHg; p=0.03), raising the possibility that the elevation in blood pressure played a role in the enhanced hemodynamic response.

**Figure 3.**
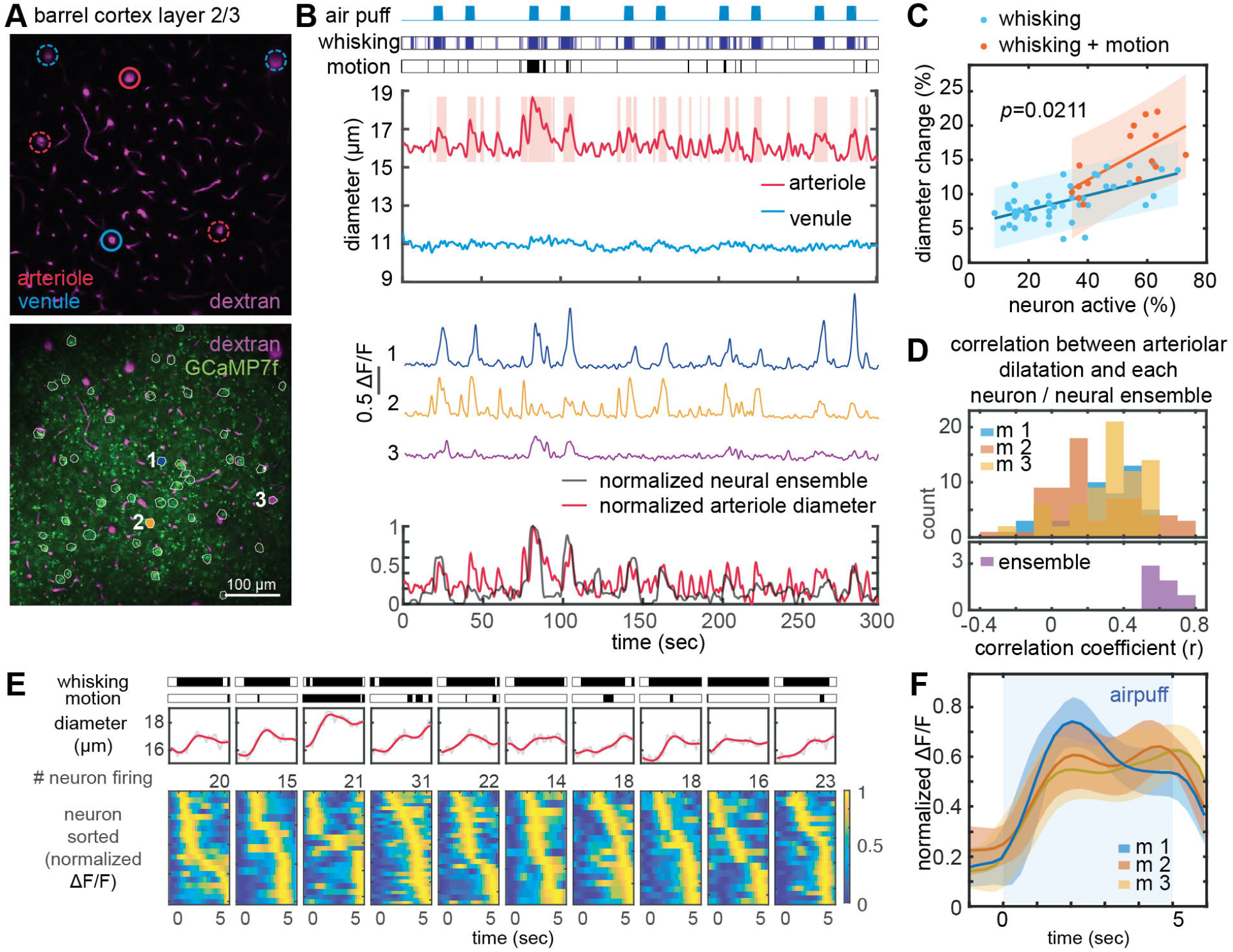
Microvascular responses associated with nNOS Ca^2+^ activity during 5 sec whisker puffs and grooming motion. **A**: 2PEF map of arterioles and venules (top) and of both vessel and GCaMP7f-expressing nNOS neurons (outlined in white, bottom). Vessels analyzed in **B** are circled in solid red or blue, and neurons are numbered (1,2,3) **B**: Arterioles, but not venules, dilate in response to whisker puffs, spontaneous whisking or motion (grooming). Representative Ca^2+^ transients in neuron 1, 2, and 3 (middle). Overlay of normalized neural ensemble activity and arteriolar diameter changes highlighting the close correspondence between arteriolar and neural responses (bottom). **C**: Linear relationship between the magnitude of the ensemble activity and arteriolar dilatation in response to whisking or whisking and spontaneous motion. Notice that with whisking+motion the slope of the relationship is significantly greater than with whisking alone (multiple comparison of one-way analysis of covariance models). **D**: The distribution of correlation coefficients between arteriolar dilatation and nNOS Ca^2+^ transients for individual neurons does not differ in the 3 mice studied (p>0.05; nested one-way ANOVA)(top). The strongest correlation is observed between ensemble activity and arteriolar dilatation (bottom). **E**: Representative arteriolar diameter changes relative to the timing of the peak Ca^2+^ activity in individual neuron during 10 consecutive 5 sec whisker air puffs shown in **B**. Notice that in some neurons peak activity occur earlier than in other neurons. **F**: Average of ten raster plots in 3 mice showing early and late active nNOS neurons. Shading indicates standard deviation.

### Arteriolar dilatation is associated with distinct phases of nNOS neuron Ca^2+^ activity

Next, we examined the relationship between individual nNOS neuron Ca^2+^ transients and arteriolar dilatation. With 5 sec air puffs, we found varying degrees of correlations (r:-0.36 to 0.77), but ensemble activity exhibited the strongest correlation with arteriolar dilatation (**Figure 3D**). We did not observe dilatation of the venules **(Figure 3B)**. The discrepancy between the lower correlation coefficient of individual neurons compared to ensemble activity raises the possibility that there are differences in the timing of neuronal activity relative to arteriolar dilatation. Therefore, we examined the temporal relationships between nNOS neuron Ca^2+^ transients and arteriolar dilatation. We found that within a single 5 second whisker air-puff, there were neurons that were active immediately as the air-puff started, and other neurons that had a delayed onset of activity (**Figure 3E, F**).

To better resolve the temporal relationships between nNOS Ca^2+^ transients and arteriolar dilatation we used a longer whisker stimulus (**Figure 4A**). With a 30 sec stimulus we observed a robust arteriolar dilatation (8.1±3.1%) associated with a small increase in venular diameter (2.0±0.8%) peaking at the end of the stimulation period (**Figure 4A**). As for the Ca^2+^ transients, a group nNOS neurons became active immediately before the arteriolar dilation (early responding neurons), their activity subsiding despite persistence of the air-puff. Other neurons (sustaining neurons) became gradually active after the dilatation and remained active throughout the air-puff (**Figure 4B,C,D**). Correlation analysis between the shape of the Ca^2+^ transient and the arteriolar dilatation revealed that the Ca^2+^ transients of the sustaining neurons were similar in shape to the arteriolar dilatation signal (**Figure 4D**), consistent with their involvement in the maintenance of the vasodilatation.

**Figure 4.**
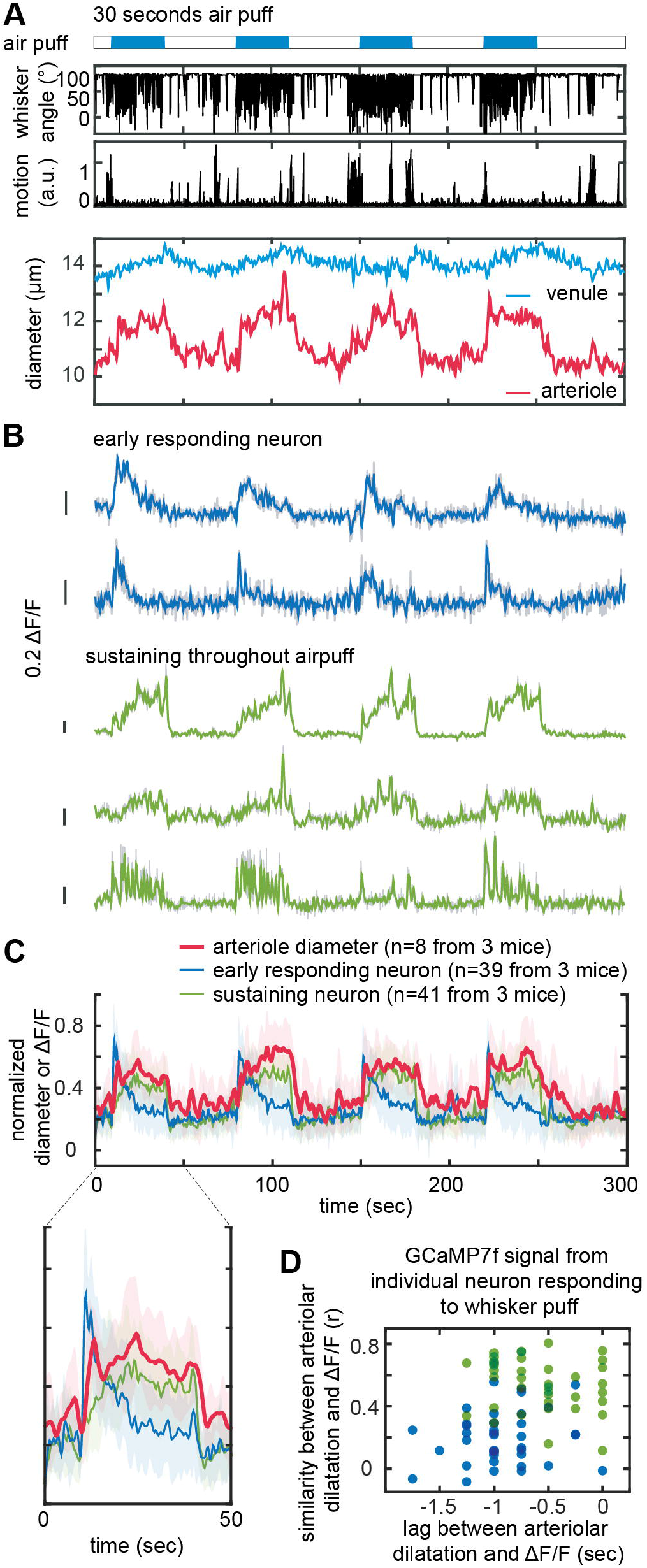
Microvascular responses associated with nNOS Ca^2+^ activity during 30 sec whisker puffs. **A**: Representative dataset from four 30-seconds whisker air puff showing the corresponding dilatation of arterioles and venules. **B**: Representative pattern of Ca^2+^ (ΔF/F) transients showing early responding neurons (blue) and neurons with sustained activity throughout the stimulus (green). **C**: Average of normalized Ca^2+^ transients and corresponding arteriolar dilatation (8 datasets in 3 mice). The enlarged graph for a single air puff shows neurons active prior to the arteriolar dilatation (early responding neurons; blue) and neurons active during the dilatation (sustaining neurons; green). **D:** plot of the time lag between Ca^2+^ changes and arteriolar dilatation, and similarity of the time course of neuronal and arteriolar responses. Two distinct nNOS neuronal populations can be identified: one rapidly decaying and preceding the arteriolar dilatation (blue; early responding neurons), and the other exhibiting a time course similar to the arteriolar dilation (green; sustaining neurons).

### The hemodynamic response is not related to the proximity of active nNOS neurons to local arterioles

nNOS neurons release NO that is thought to diffuse to nearby arterioles causing their dilatation ^25-27^. Since some nNOS neurons are closely associated with blood vessels ^21, 23, 24^, we examined whether the dilation of a particular arteriole was related to its proximity to the soma of active nNOS neurons. To this end, we calculated correlation coefficient between arteriolar dilatation and the Ca^2+^ signals for individual nNOS neurons (early responding and sustaining) in a 509×509 μm^2^ field (**Figure 5 A**). The correlation coefficient was then plotted as a function of the distance between the neurons and the arterioles. We found that there was no correlation between the proximity of an arteriole to nNOS neurons and its dilatation (**Figure 5B**, all R-Squared values close to 0).

**Figure 5.**
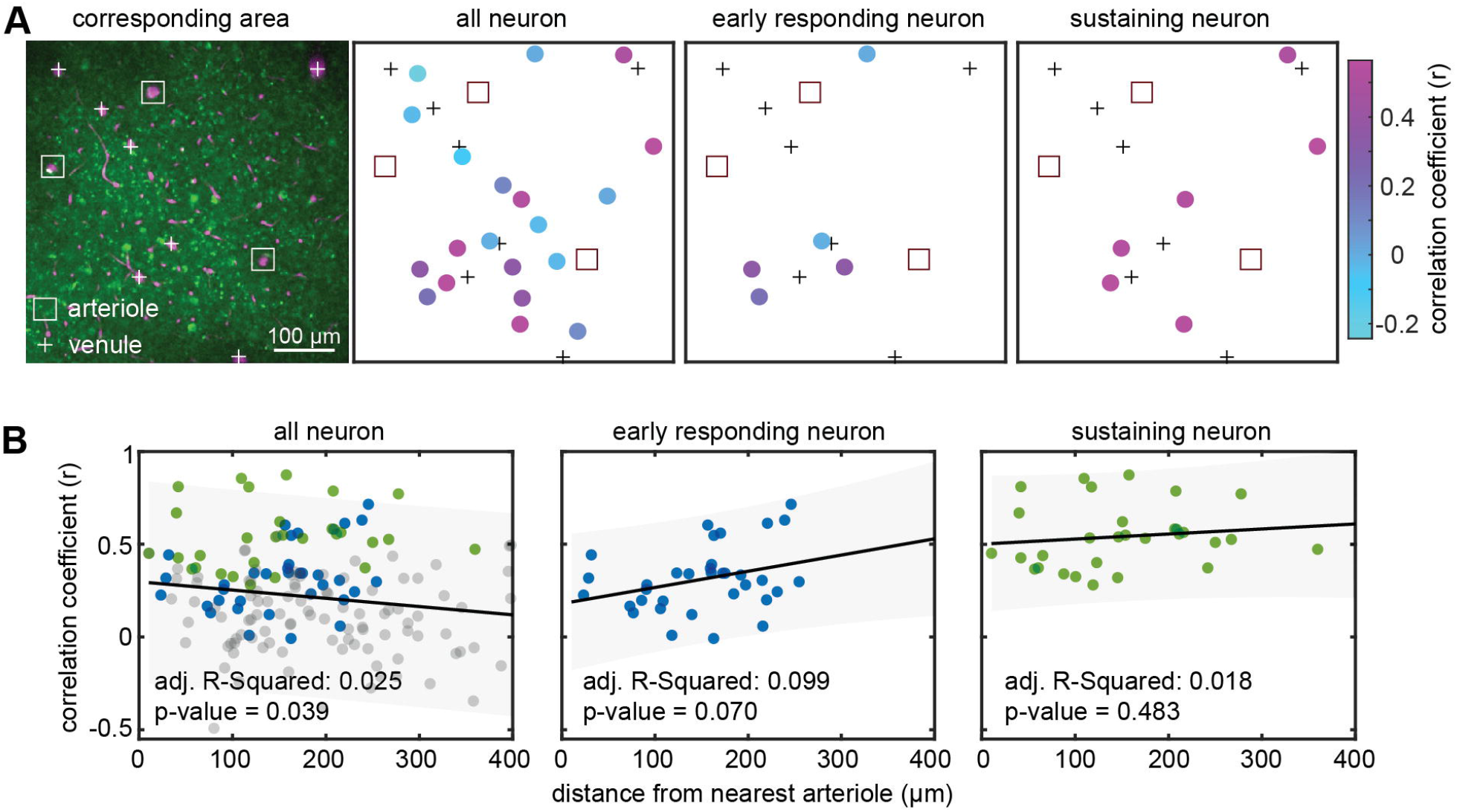
Spatial relationship of nNOS neurons responsive to air puff and arterioles. **A**: Representative 2PEF map of arterioles (square outline) and venules (+ sign) and GCaMP7f-expressing nNOS neurons. The schematic maps on the right show individual neurons color coded for the correlation of their Ca^2+^ response with that arteriolar dilatation. Notice that the highly correlated neurons do not cluster near the arterioles. **B**: Linear regression analysis between distance of individual neuron from arterioles and correlation of their Ca^2+^ response with that arteriolar dilatation (n= 170 neurons from 3 mice). There is no relationship between the distance from the nearest arteriole and correlation of Ca^2+^ transients with arteriolar dilatation.

## Discussion

We sought to investigate the relationship between nNOS neuron network activity and the corresponding microvascular changes in the somatosensory cortex of awake mice. We found that voluntary whisking, air puff-induced whisker stimulation, and grooming-related motion induced significant nNOS ensemble activity. Different patterns of Ca^2+^ transients were observed: some nNOS neurons responded to either whisking or motion, some to both, and some were active in a manner not related to these inputs. The nNOS neuron activity was associated with dilatation of the arterioles in the field of view. We noticed that the Ca^2+^ response of some nNOS neurons was closely correlated with the arteriolar dilatation but the ensemble activity of all nNOS neurons in the field of view yielded the best correlation. By investigating the timing of the Ca^2+^ signal relative to the arteriolar dilatation we discovered that some nNOS neurons became active either immediately prior to the arteriolar response or after the response was well developed, unveiling two distinct phases in the nNOS neural events associated with the vasodilation. Finally, we found that the activity of nNOS neurons closer to local arterioles was not better correlated with the hemodynamic response than that of more distant neurons, whether early or late responding. These observations provide new insights into the activation pattern of nNOS neurons, and into the relationship between nNOS neuron Ca^2+^ activity and the ensuing microvascular response.

A major strength of the present study is that the experiments were performed in awake mice, which is critically important when investigating interneuron activity, highly sensitive to anesthetics ^42^. Accordingly, we could monitor not only neurovascular coupling related to air puff-induced whisker stimulation but also to spontaneous whisking and grooming-related motion. We found that the hemodynamic response to whisking plus motion was more pronounced than that of whisking alone. However, the increase in ensemble activity could not explain the enhanced vasodilatation. We found that grooming motion was associated with an increase in blood pressure, which may also contribute since rapid changes in blood pressure escape cerebrovascular autoregulation and influence CBF ^58^. Monitoring arterial pressure and blood gasses, is challenging especially in the awake state. However, it may be needed to assess the impact of systemic factors on CBF responses, particularly in conscious mice ^2^.

We found different patterns of nNOS neuron Ca^2+^ activity in the whisker barrel cortex. Some nNOS neurons responded to either whisking or motion, some to both, and some were active independently. Such diversity is anticipated based on the complexity of the nNOS neuronal network and its connectivity ^59^. Thus, nNOS neurons are innervated by cortical and subcortical pathways forming a rich network responsive to a wide variety of stimuli ^18-20, 24, 35, 60^. In agreement with data in the literature, we found that the majority of nNOS neurons also expressed GAD67 reflecting their inhibitory nature ^13^. However, we also observed a subpopulation of nNOS neurons expressing markers of excitatory neurons, such CaMKII, a finding confirmed by mining of single-cells transcriptomic data. Since cortical nNOS neurons receive thalamic afferents ^60^, it is not surprising that some nNOS neurons responded faithfully to sensory inputs. However, we cannot comment on whether these nNOS neurons are excitatory of inhibitory or whether these are type 1 or type 2.

Another key finding of the study is that there are two distinct groups of layer 2/3 nNOS neurons linked to the hemodynamic response, one active immediately prior to the arteriolar dilatation and the other becoming active more gradually throughout the response. nNOS, a Ca^2+^-calmodulin dependent enzyme, is activated by intracellular Ca^2+ 61^. Assuming that the GCaMP7f signal reflects an increase in intracellular Ca^2+^ sufficient to activate nNOS, it is likely that the Ca^2+^ signal in these neurons is associated with NO production. Considering the wealth of data linking neuronal NO with functional hyperemia ^25-33^, it is conceivable that early active nNOS neurons are involved in the initiation of the vasodilatation. Indeed, NO release has been shown to precede the flow response during neural activation ^25-27^. As for the nNOS neurons active later, studies did not detect sustained NO release throughout the hemodynamic response ^25-27^.

However, in these studies NO readings were taken at a single point with implanted electrodes in brain slices or in vivo under anesthesia^25-27^. Furthermore, the activating stimulus, NMDA application ^25, 27^ or electrical stimulation of the forepaw ^26^, was not comparable to whisker air puffs in the awake state. Alternatively, since nNOS neurons have the potential to release also vasoactive neuropeptides ^35, 62^, we cannot rule out the possibility that the nNOS neurons responding later contribute to the response by releasing these agents. Another caveat is that 2PEF imaging performed on parallel plane to the brain surface can only visualize a small fraction of nNOS neurons in layer 2/3. Considering the heterogeneous distribution of nNOS neuron in the neocortex ^14^, we cannot exclude that other patterns of activity may be evoked by somatosensory stimuli in nNOS neurons in other layers or cortical areas.

The network events driving these two patterns of activity by different nNOS neurons remain unclear. Previous studies have revealed that the rodent whisker barrel cortex receives and processes sensory information in temporally distinct patterns. Short latency responses are mainly found in layer 4 and 5B/6, where the sensory thalamic input terminates ^63-66^. More superficial neurons in layer 2/3 respond with longer latencies and typically exhibit longer-lasting responses ^67, 68^. Furthermore, GABAergic neurons located in layer 3-5 are known to receive prominent short-latency input from the thalamus and are strongly and reciprocally connected to nearby excitatory neurons, thereby providing both feed forward and feedback inhibition within local microcircuits^69, 70^. How these complex network events engage different groups of nNOS neurons to contribute to the early and late phases of functional hyperemia remains to be determined.

We also found that the distance of the soma of nNOS neurons from arterioles did not correlate with the vascular response. This observation is somewhat unexpected since NO diffuses along a concentration gradient, and the arterioles closer to the NO source (nNOS neurons) would be exposed to a higher NO concentration. However, considering the high two-dimensional diffusion coefficient of NO (3,300 μm^2^/sec) ^71, 72^, the distance of the processes from the vessels could have no impact over the small distances examined. On the other hand, since nNOS is enriched in the terminals of nNOS neurons ^21, 23, 24^ and we monitored only the position of the soma, our spatial data needs to be interpreted with caution. Therefore, more sensitive activity markers and imaging modalities with higher resolution would be needed to provide insight into the relationship between the location of nNOS and the hemodynamic response.

In conclusion, we have demonstrated that nNOS neurons exhibit diverse patterns of activity in awake behaving mice with and without mechanical stimulation of the facial whiskers. nNOS neurons activity displayed a selective stimulus dependence to whisker stimulation or grooming and their ensemble activity was associated with robust dilatation of local arterioles. With whisker stimulation, correlation analysis revealed two distinct temporal patterns of activity one preceding and the other following the arteriolar dilatation, suggesting temporally distinct roles of nNOS neurons in the neural events underlying functional hyperemia. Interestingly, the location of nNOS neurons relative to the arteriole did not show a correlation with the hemodynamic response. Collectively, the data suggest that nNOS neurons are selectively engaged by diverse inputs and that the NO-dependent component of functional hyperemia is driven by time-dependent engagement of distinct groups of nNOS neurons. These novel findings may provide insight into the neurovascular dysfunction associated with brain diseases, such as tauopathies, amyloid pathology, and pathological stress, and in which nNOS neurons are preferentially targeted by the disease process ^44-46^.

## Acknowledgements

Supported by NIH grants R01-NS37853, R01-NS128947, R01-NS126467, and R37-NS089323 to CI and by the Feil Family Foundation. SA is a Leon Levy Neuroscience Fellow. We thank Dr. Conor Liston for assistance in the analysis of the Ca^2+^ data.

## Author contribution statement

SA, JA, and CI developed the concept and designed the experiments. SA performed the experiments and data analysis. CI and SA wrote the manuscript. CI and JA provided supervision. CI provided funding.

## References

1. Gorelick PB, Furie KL, Iadecola C, et al. Defining Optimal Brain Health in Adults: A Presidential Advisory From the American Heart Association/American Stroke Association. Stroke 2017; 48: e284–e303. 10.1161/STR.0000000000000148 2017/09/09. DOI: 10.1161/STR.0000000000000148.

2. Schaeffer S and Iadecola C. Revisiting the neurovascular unit. Nat Neurosci 2021; 24: 1198–1209. 2021/08/07. DOI: 10.1038/s41593-021-00904-7.

3. Iadecola C, Yang G, Ebner TJ, et al. Local and propagated vascular responses evoked by focal synaptic activity in cerebellar cortex. J Neurophysiol 1997; 78: 651–659. DOI: papers3://publication/uuid/C15AF186-4EFB-4BC0-9AB7-C851EF7FBCDC.

4. Longden TA, Dabertrand F, Koide M, et al. Capillary K(+)-sensing initiates retrograde hyperpolarization to increase local cerebral blood flow. Nat Neurosci 2017; 20: 717–726. 10.1038/nn.4533 2017/03/21. DOI: 10.1038/nn.4533.

5. Chen BR, Kozberg MG, Bouchard MB, et al. A critical role for the vascular endothelium in functional neurovascular coupling in the brain. J Am Heart Assoc 2014; 3: e000787. 10.1161/JAHA.114.000787 2014/06/14. DOI: 10.1161/JAHA.114.000787.

6. Thakore P, Alvarado MG, Ali S, et al. Brain endothelial cell TRPA1 channels initiate neurovascular coupling. Elife 2021; 10 2021/02/27. DOI: 10.7554/eLife.63040.

7. Attwell D and Iadecola C. The neural basis of functional brain imaging signals. Trends Neurosci 2002; 25: 621–625. DOI: 10.1016/s0166-2236(02)02264-6.

8. Silva AC and Koretsky AP. Laminar specificity of functional MRI onset times during somatosensory stimulation in rat. Proc Natl Acad Sci U S A 2002; 99: 15182–15187. 20021029. DOI: 10.1073/pnas.222561899.

9. Iadecola C. Neurovascular regulation in the normal brain and in Alzheimer’s disease. Nat Rev Neurosci 2004; 5: 347–360. 10.1038/nrn1387 2004/04/22. DOI: 10.1038/nrn1387.

10. Attwell D, Buchan AM, Charpak S, et al. Glial and neuronal control of brain blood flow. Nature 2010; 468: 232–243. 10.1038/nature09613 2010/11/12. DOI: 10.1038/nature09613.

11. Hartmann DA, Coelho-Santos V and Shih AY. Pericyte Control of Blood Flow Across Microvascular Zones in the Central Nervous System. Annu Rev Physiol 2021 20211021. DOI: 10.1146/annurev-physiol-061121-040127.

12. Hosford PS and Gourine AV. What is the key mediator of the neurovascular coupling response? Neurosci Biobehav Rev 2019; 96: 174–181. 10.1016/j.neubiorev.2018.11.011 20181124. DOI: 10.1016/j.neubiorev.2018.11.011.

13. Perrenoud Q, Geoffroy H, Gauthier B, et al. Characterization of Type I and Type II nNOS-Expressing Interneurons in the Barrel Cortex of Mouse. Front Neural Circuits 2012; 6: 36. 10.3389/fncir.2012.00036 2012/07/04. DOI: 10.3389/fncir.2012.00036.

14. Wu YT, Bennett HC, Chon U, et al. Quantitative relationship between cerebrovascular network and neuronal cell types in mice. Cell Rep 2022; 39: 110978. 2022/06/23. DOI: 10.1016/j.celrep.2022.110978.

15. Taniguchi H. Genetic dissection of GABAergic neural circuits in mouse neocortex. Front Cell Neurosci 2014; 8: 8. 2014/01/31. DOI: 10.3389/fncel.2014.00008.

16. Huang ZJ and Paul A. The diversity of GABAergic neurons and neural communication elements. Nat Rev Neurosci 2019; 20: 563–572. 2019/06/22. DOI: 10.1038/s41583-019-0195-4.

17. Lourenço CF and Laranjinha J. Nitric Oxide Pathways in Neurovascular Coupling Under Normal and Stress Conditions in the Brain: Strategies to Rescue Aberrant Coupling and Improve Cerebral Blood Flow. Frontiers in Physiology 2021; 12. DOI: 10.3389/fphys.2021.729201.

18. Gerashchenko D, Wisor JP, Burns D, et al. Identification of a population of sleep-active cerebral cortex neurons. Proc Natl Acad Sci U S A 2008; 105: 10227–10232. 20080721. DOI: 10.1073/pnas.0803125105.

19. Gerashchenko D, Wisor JP and Kilduff TS. Sleep-active cells in the cerebral cortex and their role in slow-wave activity. Sleep Biol Rhythms 2011; 9: 71–77. 2011/06/01. DOI: 10.1111/j.1479-8425.2010.00461.x.

20. Morairty SR, Dittrich L, Pasumarthi RK, et al. A role for cortical nNOS/NK1 neurons in coupling homeostatic sleep drive to EEG slow wave activity. Proc Natl Acad Sci U S A 2013; 110: 20272–20277. 20131104. DOI: 10.1073/pnas.1314762110.

21. Wang H, Hitron IM, Iadecola C, et al. Synaptic and vascular associations of neurons containing cyclooxygenase-2 and nitric oxide synthase in rat somatosensory cortex. Cereb Cortex 2005; 15: 1250–1260. 10.1093/cercor/bhi008 2004/12/24. DOI: 10.1093/cercor/bhi008.

22. Duchemin S, Boily M, Sadekova N, et al. The complex contribution of NOS interneurons in the physiology of cerebrovascular regulation. Front Neural Circuits 2012; 6: 51. 2012/08/22. DOI: 10.3389/fncir.2012.00051.

23. Iadecola C, Beitz AJ, Renno W, et al. Nitric oxide synthase-containing neural processes on large cerebral arteries and cerebral microvessels. Brain Res 1993; 606: 148–155. 1993/03/19. DOI: 10.1016/0006-8993(93)91583-e.

24. Tong X and Hamel E. Basal forebrain nitric oxide synthase (NOS)lJcontaining neurons project to microvessels and NOS neurons in the rat neocortex-cellular basis for cortical blood flow regulation.pdf. european journal of neuroscince 2020.

25. Rancillac A, Rossier J, Guille M, et al. Glutamatergic Control of Microvascular Tone by Distinct GABA Neurons in the Cerebellum. J Neurosci 2006; 26: 6997–7006. 10.1523/JNEUROSCI.5515-05.2006 2006/06/30. DOI: 10.1523/JNEUROSCI.5515-05.2006.

26. Buerk DG, Ances BM, Greenberg JH, et al. Temporal dynamics of brain tissue nitric oxide during functional forepaw stimulation in rats. Neuroimage 2003; 18: 1–9. 2003/01/01. DOI: 10.1006/nimg.2002.1314.

27. Lourenco CF, Santos RM, Barbosa RM, et al. Neurovascular coupling in hippocampus is mediated via diffusion by neuronal-derived nitric oxide. Free Radic Biol Med 2014; 73: 421–429. 10.1016/j.freeradbiomed.2014.05.021 2014/06/03. DOI: 10.1016/j.freeradbiomed.2014.05.021.

28. Yang G, Huard JM, Beitz AJ, et al. Stellate neurons mediate functional hyperemia in the cerebellar molecular layer. J Neurosci 2000; 20: 6968-6973. 2000/09/21. DOI: papers3://publication/uuid/AB196B4A-BABA-436D-A9C3-4D6DE331F490.

29. Yang G and Iadecola C. Obligatory role of NO in glutamate-dependent hyperemia evoked from cerebellar parallel fibers. Am J Physiol 1997; 272: R1155–1161. DOI: papers3://publication/uuid/28BD1077-62AF-44E3-8689-FC10AAB8EE03.

30. Faraci F and Brian J. 7-Nitroindazole inhibits brain nitric oxide synthase and cerebral vasodilatation in response to N-methyl-D-aspartate. Stroke 1995; 26: 2172–2176. DOI: papers3://publication/uuid/12B99A59-3B7F-4602-9A86-26905066AD5D.

31. Bonvento G, Cholet N and Seylaz J. Sustained attenuation of the cerebrovascular response to a 10 min whisker stimulation following neuronal nitric oxide synthase inhibition. Neurosci Res 2000; 37: 163–166. DOI: papers3://publication/uuid/0B7C120D-194B-4BE4-960C-9811F3778E63.

32. Liu X, Li C, Falck JR, et al. Interaction of nitric oxide, 20-HETE, and EETs during functional hyperemia in whisker barrel cortex. Am J Physiol Heart Circ Physiol 2008; 295: H619–631. 2008/05/27. DOI: 10.1152/ajpheart.01211.2007.

33. Vazquez AL, Fukuda M and Kim SG. Inhibitory Neuron Activity Contributions to Hemodynamic Responses and Metabolic Load Examined Using an Inhibitory Optogenetic Mouse Model. Cereb Cortex 2018; 28: 4105–4119. 10.1093/cercor/bhy225 2018/09/15. DOI: 10.1093/cercor/bhy225.

34. Hoiland RL, Caldwell HG, Howe CA, et al. Nitric oxide is fundamental to neurovascular coupling in humans. J Physiol (Lond) 2020; 598: 4927–4939. 10.1113/JP280162 2020/08/14. DOI: 10.1113/JP280162.

35. Cauli B, Tong XK, Rancillac A, et al. Cortical GABA interneurons in neurovascular coupling: relays for subcortical vasoactive pathways. J Neurosci 2004; 24: 8940–8949. 10.1523/JNEUROSCI.3065-04.2004 2004/10/16. DOI: 10.1523/JNEUROSCI.3065-04.2004.

36. Echagarruga CT, Gheres KW, Norwood JN, et al. nNOS-expressing interneurons control basal and behaviorally evoked arterial dilation in somatosensory cortex of mice. Elife 2020; 9 2020/10/06. DOI: 10.7554/eLife.60533.

37. Lee L, Boorman L, Glendenning E, et al. Key Aspects of Neurovascular Control Mediated by Specific Populations of Inhibitory Cortical Interneurons. Cereb Cortex 2020; 30: 2452-2464. 2019/11/21. DOI: 10.1093/cercor/bhz251.

38. Krawchuk MB, Ruff CF, Yang X, et al. Optogenetic assessment of VIP, PV, SOM and NOS inhibitory neuron activity and cerebral blood flow regulation in mouse somato-sensory cortex. J Cereb Blood Flow Metab 2020; 40: 1427–1440. 2019/08/17. DOI: 10.1177/0271678X19870105.

39. Slupe AM and Kirsch JR. Effects of anesthesia on cerebral blood flow, metabolism, and neuroprotection. J Cereb Blood Flow Metab 2018; 38: 2192–2208. 2018/07/17. DOI: 10.1177/0271678X18789273.

40. Franceschini MA, Radhakrishnan H, Thakur K, et al. The effect of different anesthetics on neurovascular coupling. Neuroimage 2010; 51: 1367–1377. 2010/03/31. DOI: 10.1016/j.neuroimage.2010.03.060.

41. Masamoto K and Kanno I. Anesthesia and the quantitative evaluation of neurovascular coupling. J Cereb Blood Flow Metab 2012; 32: 1233–1247. 10.1038/jcbfm.2012.50 2012/04/19. DOI: 10.1038/jcbfm.2012.50.

42. Guo J, Ran M, Gao Z, et al. Cell-type-specific imaging of neurotransmission reveals a disrupted excitatory-inhibitory cortical network in isoflurane anaesthesia. EBioMedicine 2021; 65: 103272. 2021/03/11. DOI: 10.1016/j.ebiom.2021.103272.

43. Gao YR, Ma Y, Zhang Q, et al. Time to wake up: Studying neurovascular coupling and brain-wide circuit function in the un-anesthetized animal. Neuroimage 2017; 153: 382–398. 10.1016/j.neuroimage.2016.11.069 2016/12/03. DOI: 10.1016/j.neuroimage.2016.11.069.

44. Park L, Hochrainer K, Hattori Y, et al. Tau induces PSD95-neuronal NOS uncoupling and neurovascular dysfunction independent of neurodegeneration. Nat Neurosci 2020; 23: 1079–1089. 10.1038/s41593-020-0686-7 2020/08/12. DOI: 10.1038/s41593-020-0686-7.

45. Choi S, Won JS, Carroll SL, et al. Pathology of nNOS-Expressing GABAergic Neurons in Mouse Model of Alzheimer’s Disease. Neuroscience 2018; 384: 41–53. 2018/05/22. DOI: 10.1016/j.neuroscience.2018.05.013.

46. Han K, Min J, Lee M, et al. Neurovascular Coupling under Chronic Stress Is Modified by Altered GABAergic Interneuron Activity. J Neurosci 2019; 39: 10081–10095. 2019/11/02. DOI: 10.1523/JNEUROSCI.1357-19.2019.

47. Percie du Sert N, Hurst V, Ahluwalia A, et al. The ARRIVE guidelines 2.0: updated guidelines for reporting animal research. J Physiol (Lond) 2020; 598: 3793–3801. 10.1113/JP280389 2020/07/16. DOI: 10.1113/JP280389.

48. Ahn SJ, Anrather J, Nishimura N, et al. Diverse Inflammatory Response After Cerebral Microbleeds Includes Coordinated Microglial Migration and Proliferation. Stroke 2018; 49: 1719–1726. 2018/05/31. DOI: 10.1161/STROKEAHA.117.020461.

49. Niwa K, Younkin L, Ebeling C, et al. Abeta 1-40-related reduction in functional hyperemia in mouse neocortex during somatosensory activation. Proc Natl Acad Sci U S A 2000; 97: 9735–9740. DOI: papers3://publication/uuid/727C41DD-52FF-4F13-A44C-3DCD10468050.

50. Pnevmatikakis EA and Giovannucci A. NoRMCorre: An online algorithm for piecewise rigid motion correction of calcium imaging data. J Neurosci Methods 2017; 291: 83–94. 2017/08/08. DOI: 10.1016/j.jneumeth.2017.07.031.

51. Ackman JB, Burbridge TJ and Crair MC. Retinal waves coordinate patterned activity throughout the developing visual system. Nature 2012; 490: 219–225. DOI: 10.1038/nature11529.

52. Moda-Sava RN, Murdock MH, Parekh PK, et al. Sustained rescue of prefrontal circuit dysfunction by antidepressant-induced spine formation. Science 2019; 364: 1–13. 10.1126/science.aat8078 2019/04/13. DOI: 10.1126/science.aat8078.

53. Gao YR and Drew PJ. Determination of vessel cross-sectional area by thresholding in Radon space. J Cereb Blood Flow Metab 2014; 34: 1180–1187. 20140416. DOI: 10.1038/jcbfm.2014.67.

54. Winder AT, Echagarruga C, Zhang Q, et al. Weak correlations between hemodynamic signals and ongoing neural activity during the resting state. Nat Neurosci 2017; 20: 1761–1769. 10.1038/s41593-017-0007-y 2017/12/01. DOI: 10.1038/s41593-017-0007-y.

55. Garcia-Bonilla L, Sciortino R, Shahanoor Z, et al. Role of microglial and endothelial CD36 in post-ischemic inflammasome activation and interleukin-1beta-induced endothelial activation. Brain Behav Immun 2021; 95: 489–501. 2021/04/20. DOI: 10.1016/j.bbi.2021.04.010.

56. La Manno G, Siletti K, Furlan A, et al. Molecular architecture of the developing mouse brain. Nature 2021 2021/07/30. DOI: 10.1038/s41586-021-03775-x.

57. Stuart T, Butler A, Hoffman P, et al. Comprehensive Integration of Single-Cell Data. Cell 2019; 177: 1888–1902 e1821. 20190606. DOI: 10.1016/j.cell.2019.05.031.

58. Claassen J, Thijssen DHJ, Panerai RB, et al. Regulation of cerebral blood flow in humans: physiology and clinical implications of autoregulation. Physiol Rev 2021; 101: 1487–1559. 2021/03/27. DOI: 10.1152/physrev.00022.2020.

59. Cauli B, Kubota Y and Tricoire L. Cortical NO interneurons: from embryogenesis to functions. Frontiers E-books, 2014.

60. Okoro SU, Goz RU, Njeri BW, et al. Organization of cortical and thalamic input to inhibitory neurons in mouse motor cortex. bioRxiv 2022. DOI: 10.1101/2021.07.08.451716.

61. Bredt D and Snyder S. Isolation of nitric oxide synthetase, a calmodulin-requiring enzyme. Proc Natl Acad Sci U S A 1990; 87: 682–685. DOI: papers3://publication/uuid/C99FD017-103C-4B1D-8FE1-CC93C8A17949.

62. Uhlirova H, Kilic K, Tian P, et al. Cell type specificity of neurovascular coupling in cerebral cortex. Elife 2016; 5: 155. 10.7554/eLife.14315 20160531. DOI: 10.7554/eLife.14315.

63. Zhu JJ and Connors BW. Intrinsic firing patterns and whisker-evoked synaptic responses of neurons in the rat barrel cortex. J Neurophysiol 1999; 81: 1171–1183. 1999/03/23. DOI: 10.1152/jn.1999.81.3.1171.

64. Moore CI and Nelson SB. Spatio-temporal subthreshold receptive fields in the vibrissa representation of rat primary somatosensory cortex. J Neurophysiol 1998; 80: 2882–2892. 1998/12/24. DOI: 10.1152/jn.1998.80.6.2882.

65. Constantinople CM and Bruno RM. Deep cortical layers are activated directly by thalamus. Science 2013; 340: 1591–1594. 2013/07/03. DOI: 10.1126/science.1236425.

66. El-Boustani S, Sermet BS, Foustoukos G, et al. Anatomically and functionally distinct thalamocortical inputs to primary and secondary mouse whisker somatosensory cortices. Nat Commun 2020; 11: 3342. 20200703. DOI: 10.1038/s41467-020-17087-7.

67. Crochet S, Poulet JF, Kremer Y, et al. Synaptic mechanisms underlying sparse coding of active touch. Neuron 2011; 69: 1160–1175. 2011/03/26. DOI: 10.1016/j.neuron.2011.02.022.

68. Zhang W and Bruno RM. High-order thalamic inputs to primary somatosensory cortex are stronger and longer lasting than cortical inputs. Elife 2019; 8 20190211. DOI: 10.7554/eLife.44158.

69. Gabernet L, Jadhav SP, Feldman DE, et al. Somatosensory integration controlled by dynamic thalamocortical feed-forward inhibition. Neuron 2005; 48: 315–327. 2005/10/26. DOI: 10.1016/j.neuron.2005.09.022.

70. Cruikshank SJ, Lewis TJ and Connors BW. Synaptic basis for intense thalamocortical activation of feedforward inhibitory cells in neocortex. Nat Neurosci 2007; 10: 462–468. 2007/03/06. DOI: 10.1038/nn1861.

71. Ledo A, Barbosa R, Gerhardt G, et al. Concentration dynamics of nitric oxide in rat hippocampal subregions evoked by stimulation of the NMDA glutamate receptor. Proc Natl Acad Sci U S A 2005; 102: 17483–17488. DOI: papers3://publication/uuid/607C6AB1-3C13-4EDF-9002-A3D93D4D7219.

72. Lancaster JR, Jr. A tutorial on the diffusibility and reactivity of free nitric oxide. Nitric Oxide 1997; 1: 18–30. DOI: 10.1006/niox.1996.0112.

